# A blueprint of seed desiccation sensitivity in the genome of *Castanospermum australe*

**DOI:** 10.1101/665661

**Authors:** Alexandre Marques, Maria-Cecília D. Costa, Udisha Chathuri, Eef Jonkheer, Tao Zhao, Elio Schijlen, Martijn Derks, Harm Nijveen, Marina Marcet-Houben, Irene Julca, Julien Delahaie, M. Eric Schranz, Toni Gabaldon, Sandra Pelletier, Olivier Leprince, Wilco Ligterink, Julia Buitink, Henk W.M. Hilhorst, Jill M. Farrant

## Abstract

- Most angiosperms produce seeds that are desiccated on dispersal with the ability to retain viability in storage facilities for prolonged periods. However, some species produce desiccation sensitive seeds which rapidly lose viability in storage, precluding *ex situ* conservation. Current consensus is that desiccation sensitive seeds either lack or do not express mechanisms necessary for the acquisition of desiccation tolerance.
- We sequenced the genome of *Castanospermum australe*, a legume species producing desiccation sensitive seeds, and characterized its seed developmental physiology and - transcriptomes.
- *C. australe* has a low rate of evolution, likely due to its perennial life-cycle and long generation times. The genome is syntenic with itself, with several orthologs of genes from desiccation tolerant legume seeds, from gamma whole-genome duplication events being retained. Changes in gene expression during development of *C. australe* seeds, as compared to desiccation tolerant *Medicago truncatula* seeds, suggest they remain metabolically active, prepared for immediate germination.
- Our data indicates that the phenotype of *C. australe* seeds arose through few changes in specific signalling pathways, precluding or bypassing activation of mechanisms necessary for acquisition of desiccation tolerance. Such changes have been perpetuated as the habitat in which dispersal occurs is favourable for prompt germination.

## Introduction

Seeds of most gymnosperm and angiosperm species are shed in the desiccated state and can be stored dry under sub-zero temperatures for prolonged periods of time, thus facilitating plant germplasm conservation. Such desiccation tolerant (DT) seeds are termed ‘orthodox’. However, seeds of some species are desiccation-sensitive (DS), also referred to as ‘recalcitrant’, and cannot be successfully stored under typical conditions (Berjak & Pammenter, 2013). DS-seeded species are mostly found in the humid tropics and may represent up to 50% of the species present in tropical evergreen rain forests (Hamilton *et al.*, 2013). Such species occur in environments conducive to immediate seed germination and thus selective pressure for desiccation tolerance has been relaxed or is absent. It has been hypothesized that seed desiccation sensitivity is a derived trait that evolved independently in non-related clades (Berjak & Pammenter, 2000). Genes responsible for seed desiccation tolerance would have been lost, repressed and/or mutated in DS seeded species (Berjak & Pammenter, 2008). However, this hypothesis remains to be tested at the genome level. Here we applied an extensive phylogenetic comparison to obtain a genomic blueprint of desiccation sensitivity in seeds.

Desiccation tolerance is acquired mid-way during the development of orthodox seeds when seed filling is approximately half-way, corresponding to a steep drop in water content of the seeds, concomitantly with a transient rise in abscisic acid (ABA) content (Bewley *et al.*, 2013). This acquisition comprises highly coordinated molecular events, including the repression of photosynthesis and energy metabolism, and accumulation of protective components, such as late embryogenesis abundant (LEA) proteins, anti-oxidants, and soluble sugars (Leprince *et al.*, 2017). These events are tightly regulated by hormones such as ABA and transcription factors (TFs) such as *ABSCISIC ACID INSENSITIVE 3* (*ABI3*), *FUSCA 3* (*FUS3*) and *LEAFY COTYLEDON 1* (*LEC1*) (Leprince *et al.*, 2017). Conversely, in the development of DS seeds, the acquisition of desiccation tolerance and accumulation of the above-mentioned protectants appear to be suppressed and rather they directly progress towards germination (Farrant *et al.*, 1993b; Francini *et al.*, 2006; Delahaie *et al.*, 2013). However, the genetic makeup underlying the DS seed phenotype is unknown.

The legume family (Fabaceae) contains many agriculturally important species, all producing DT seeds, such as soybean (*Glycine max*), the common bean (*Phaseolus vulgaris*), lentils (*Lens culinaris*) and chickpeas (*Cicer arietinum*). Moreover, legume species, such as soybean and *Medicago truncatula*, are important experimental models for molecular and physiological studies on seed desiccation tolerance (Chatelain *et al.*, 2012; Delahaie *et al.*, 2013; Verdier *et al.*, 2013; Zinsmeister *et al.*, 2016). In addition, the genomes of 16 DT-seeded legume species have been sequenced. Thus, a large amount of information is available, allowing comparative analysis among them.

*Castanopermum australe* A. Cunn & C. Fraser ex Hook, also known as the Moreton Bay Chestnut or Blackbean, is a leguminous tropical tree native to the east coast of Australia and west Pacific islands. In contrast to most legume species, *C. australe* produces DS seeds. It is the only known species in the genus that forms a separate and early branching clade within the Papilionoideae subfamily (Cardoso *et al.*, 2012). The earliest-branching papilionoids fall within an ADA clade, which includes the monophyletic tribes Angylocalyceae, Dipterygeae, and Amburanae. *C. australe*’s important phylogenetic position in the basis of the ADA clade (Angylocalyaceae) makes its genome ideal to study both trait evolution and the ancient polyploid history of papilionoid legumes (Schranz *et al.*, 2012).

Here, we provide detailed genomic sequence information of *C. australe* combined with time-resolved gene expression analysis of seed development of this species, including the comparison with other species producing either DS or DT seeds. Such information is key to understanding mechanisms of desiccation tolerance and, ultimately, to design strategies to improve tolerance of extreme water loss in DS seeds for conservation purposes. We investigated genomic changes associated with seed desiccation sensitivity, including gene deletions, severe mutations and gene mis-expression, as well as their relationship with gene expression patterns during seed development and maturation.

## Materials and Methods

### Plant material

A population of trees of *C. australe* growing in Pietermaritzburg (Kwazulu-Natal Province, South Africa) was the source of plant material for this work. Seed development occurs over a 6-month period, during which pods were harvested weekly. Seeds were extracted and, following histodifferentiation, were separated into component tissues (axis, cotyledons and seed coat). The following was determined annually over 2 seasons. Whole seed mass and that of component tissues (n = 60) and water content (n=10-20) was determined gravimetrically by oven drying. The ability of intact seeds to germinate was tested by planting in vermiculite. Once germinable, the amount of water loss tolerated by axes and cotyledons was determined by flash drying (Berjak *et al.*, 1990). Axis survival was determined as the ability to produce both shoots and roots when cultured in vitro on full strength MS medium. Survival of cotyledons was assessed by tetrazolium staining followed by spectrometric analysis (Sershen *et al.*, 2012). Critical water contents (calculated on a gH_2_O.g^-1^ dry mass) were calculated at those stages at which 50% survival was observed.

### Sugar and ABA content and determination

Sugars were extracted from frozen and lyophilised seeds and analysed by HPLC on a Carbopac PA-1 column (Rosnoblet *et al.*, 2007) (Dionex Corp., Sunnyvale, CA, USA). Three independent extractions and assays were performed on approx. 100 mg of tissue.

ABA was extracted and quantified as described by (Floková *et al.*, 2014).

### Genome sequencing and assembly

Freeze-dried leaf material was used for DNA isolation as described by (Bernatzky & Tanksley, 1986) with modifications. The genomic *C. australe* library consisted of 30x coverage PacBio with a mean read length of 7.8 Kb. In addition, an Illumina paired end library with reads of 100 bp and a 200-400 bp insert size was constructed and sequenced to a 64x coverage. Reads originating for contaminants were removed from all sequence data prior to assembly. Organelle genomes were also removed from the main genome assembly. Illumina reads were error-corrected using Lighter (Song *et al.*, 2014) and assembled using SparseAssembler (Ye *et al.*, 2012). A hybrid assembly was produced with DBG2OLC (Ye *et al.*, 2016) and the contigs were reordered and connected into scaffolds using SSPACE-LongRead (Boetzer & Pirovano, 2014). The assembly was polished using Sparc (Ye *et al.*, 2012) and Pilon (Walker *et al.*, 2014). PBJelly2 (English *et al.*, 2012) was used for gap closure and genome improvement. Alignments due to gene duplication and repeats were filtered out using the delta-filter utility of the MUMmer package (Kurtz *et al.*, 2004). The assembly was validated by mapping the available RNA and DNA libraries to the genome with Bowtie2 (Langmead & Salzberg, 2012) and Blasr (Chaisson & Tesler, 2012). Assembly statistics were calculated using QUAST (Gurevich *et al.*, 2013). Gene space completeness was measured using BUSCO (Benchmarking Universal Single-Copy Orthologs (Simão *et al.*, 2015)). The MAKER2 annotation pipeline (Holt & Yandell, 2011) was applied for gene prediction and repeat annotation.

### Transcriptome analysis

*C. australe* seeds were harvested at weekly intervals after seed set and at shedding from trees growing in Pietermaritzburg in 2009 and 2011. Six developmental stages were collected for cotyledon tissues, and three stages for the embryonic axes. Prior to maturation, when embryos were small, cotyledon tissue was used. For the green pod stage (Figure 2), axes and cotyledons were separated. Transcriptome analysis was performed on newly designed 12×135K Nimblegen arrays (IRHS_Ca_102K_v1) for *C. australe*, based on the genome assembly of (Delahaie *et al.*, 2013). RNA amplification, labelling and hybridization was performed according to (Terrasson *et al.*, 2013). Four biological replicates were analysed per developmental stage using the dye-swap method, and statistical analysis on the gene expression data was performed according to (Verdier *et al.*, 2013). Data are deposited in the NCBI Gene Expression Omnibus database (accession no. GSE109217, samples GSM2935461-GSM2935508). A gene was considered differentially expressed if P ≤ 0.05 in at least one comparison (axis or cotyledon) after the application of linear modelling.

**Figure 1.**
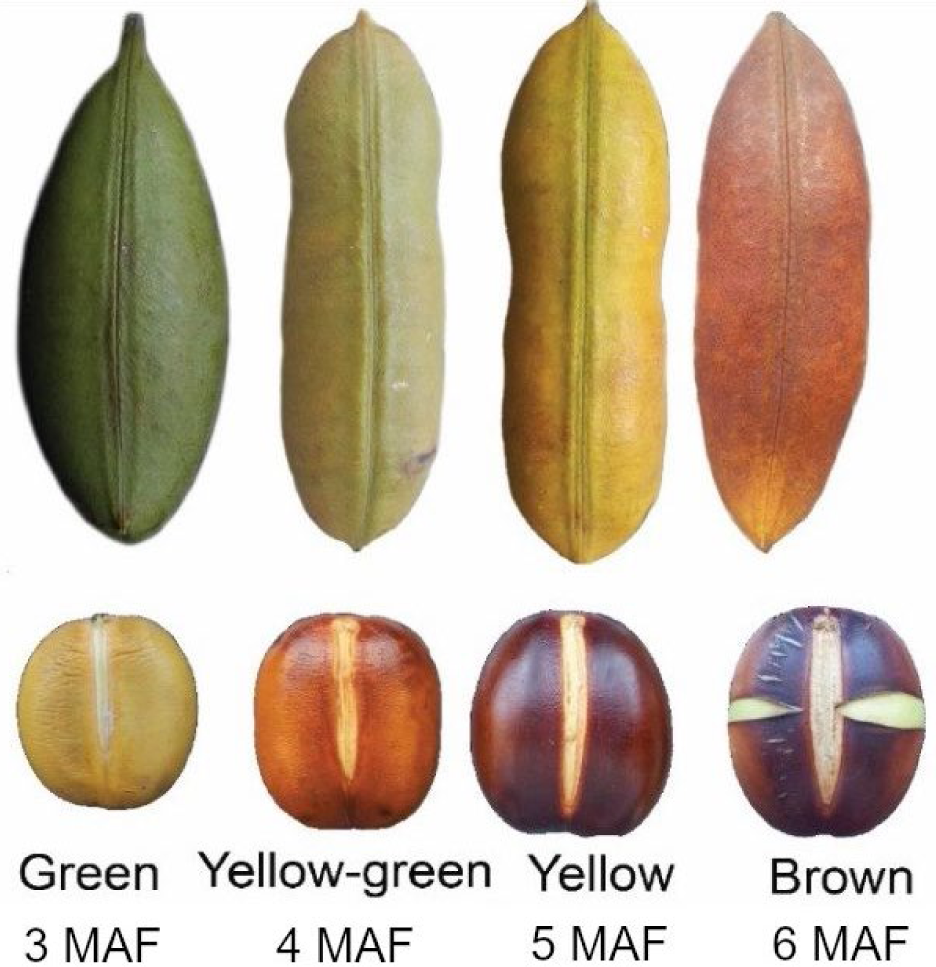
*Castanospermum australe* seed and pod developmental stages. MAF: months after flowering.

**Figure 2.**
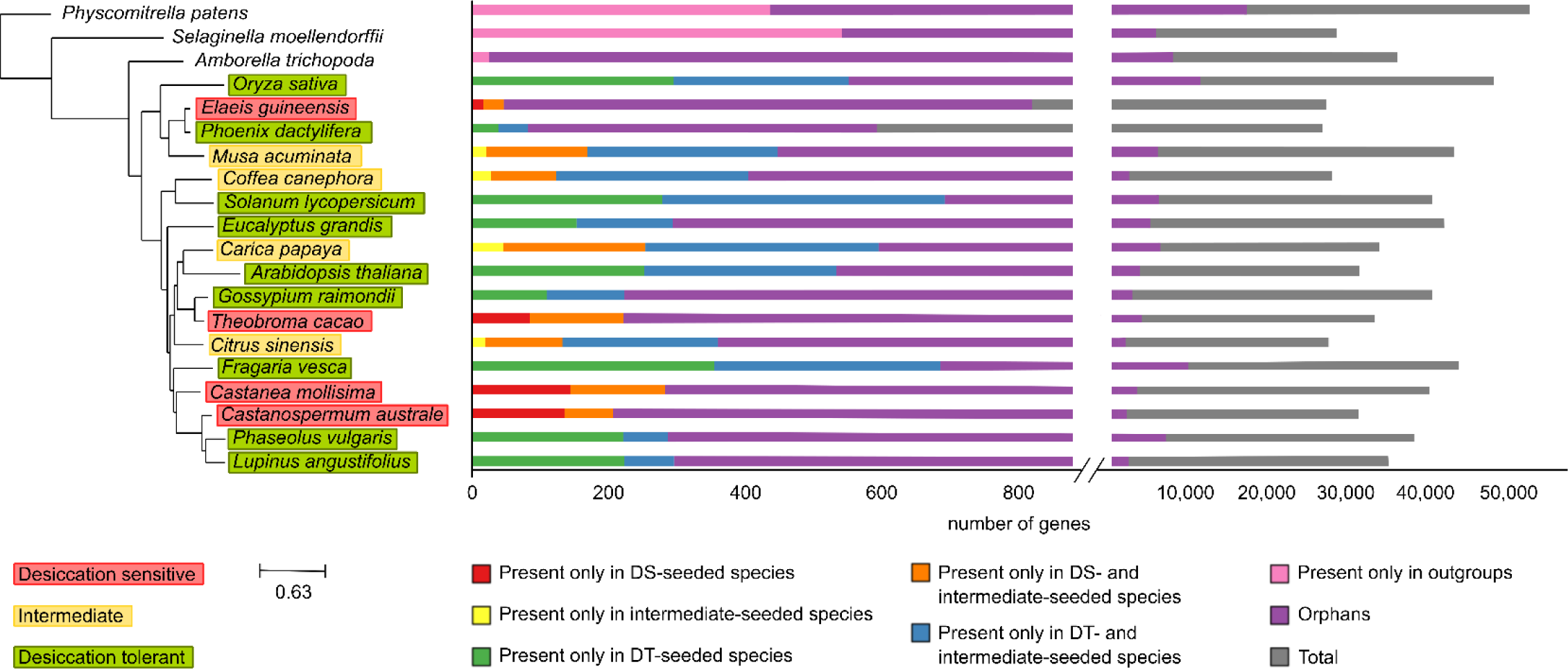
Phylome. Reconstruction of the phylome was based on a concatenated alignment of 183 single copy proteins that are present in at least 19 out of the 20 species surveyed. Species names are coloured according to their seed storage category.

Over-representation analysis (ORA) was used to recover over-represented biological processes using the app BiNGO (default settings) (Maere *et al.*, 2005) for Cytoscape.

Orthologs were defined as hits with lowest Expect value (E-value) observing a threshold of <10^-10^. Multiple hits were considered orthologs when the difference between their E-values and the lowest hit’s E-value was smaller than 10^-10^.

Members of the eight LEA protein families were identified uploading Hidden Markov Models (HMM) for each family from the PFAM database (Finn *et al.*, 2011) to HMMER 3.1b2 package (Eddy, 2011). All proteins with significant hits (E-value ≤ 0.01) were selected.

### Phylome reconstruction

We reconstructed three phylomes, one for a species set closely related to *C. australe* (phylome 110) and starting in *C. australe*, a second one based on a broader taxonomic focus also starting in *C. australe* (111) and a third one also broad but starting in *Fragaria vesca* (112). The phylome 112 was used to search for lost genes in *C. australe.* It starts in *F. vesca* that is the closest outgroup DT seeded species with a high-quality genome sequence. Both phylomes were reconstructed using the same approach (Huerta-Cepas & Gabaldon, 2011). All data generated during the phylome reconstruction has been deposited in phylomeDB (PMID:24275491) under the phylomeID codes 110 and 111. The trees, alignments and orthology and paralogy predictions are accessible to browse or download at the PhylomeDB database (Huerta-Cepas *et al.*, 2014). A set of 183 one-to-one orthologous proteins present across the compared species was used to reconstruct a species phylogeny.

### Whole genome duplication analysis

The *C. australe* genome was compared to genomic data from five Legumes: *Glycine max, Lotus japonicus, Medicago truncatula, Phaseolus vulgaris, Trifolium pratense. G. max* and *M. truncatula* are tetraploids and the rest are diploid species. The diploid *Fragaria vesca* species, of the family Rosaceae, was selected as outgroup as it is one of the closest relatives with a completed genome available outside of the Legume family. The assemblies of: *G. max, L. japonicus*, and *M. truncatula* are on chromosome level which made it easier to identify genome collinearity and duplication patterns.

### Synteny analysis

A synteny network approach (Zhao *et al.*, 2017) was implemented to compare the synteny of *C. australe* and other whole-genome sequenced legume species available at the Legume Information System (https://legumeinfo.org). BLASTP was used for pairwise genome comparison. MCScanX (Wang *et al.*, 2012) was used for synteny block detection. Infomap algorithm (Rosvall & Bergstrom, 2008) implemented in R igraph package was used for synteny network clustering. Clusters containing genes/nodes from more than 8 (out of the 10) legume species but no *C. australe* node(s) were screened out for further investigation. The maximum distance between two matches was 20 genes, a syntenic block consists of minimum 5 genes and no blocks were merged. Quota Align was enabled to determine the syntenic depth (the number of times a genomic region is syntenic). For calculating the fractionation bias, the window size was lowered from 100 to 25 considering the smaller contigs of the *C. australe* genome. Synonymous (Ks) and non-synonymous (Kn) site mutations were calculated for each syntenic gene pair. Mutation rates were used to determine if genes were duplicated by the WGD event or not. The distribution of the Ks rate was used to set a different cut-off per species.

### dN/dS analysis

We used the genome of 5 legume species from phytozome (*M. truncatula, G. max, Glycine soja, T. pratense* and *P. vulgaris*) and *C. australe*. The SynMap tool in the online CoGe portal was used to find syntenic gene pairs within these species. We set the DAGChainer on a maximum distance between two matches for 50 genes and minimum number of aligned pairs on 3 genes. To establish sets of orthologous among the 5 species against *M. truncatula*, the method of reciprocal best hits using Last was used.

Codeml in the Phylogenetic Analysis by Maximum Likelihood (PAML) package was used to estimate the dN (the rate of non-synonymous substitutions), dS (the rate of synonymous substitutions) and the ratio of dN/dS (Yang, 2007). Orthologs with dS>5, dN>2 or dN/dS>2 were filtered. For genes with multiple syntelogs we kept the pair with the lowest dN/dS.

## Results

### *Castanospermum australe* seed development

Seed development in *C. australe* occurs over a period of 6 months, with reserve accumulation and maximum embryo size completed approximately 3 months after flowering and coincident with pods becoming yellow (Figure 1 and Table 1). Unlike the embryo, the seed coat declined in mass until just prior to the yellow-green pod stage, with no further decline once embryos reached full size. Water content declined in all tissues once reserve accumulation was complete at the yellow pod stage. This loss stabilized in all tissues but the seed coat, which continued to lose water.

**Table 1.**
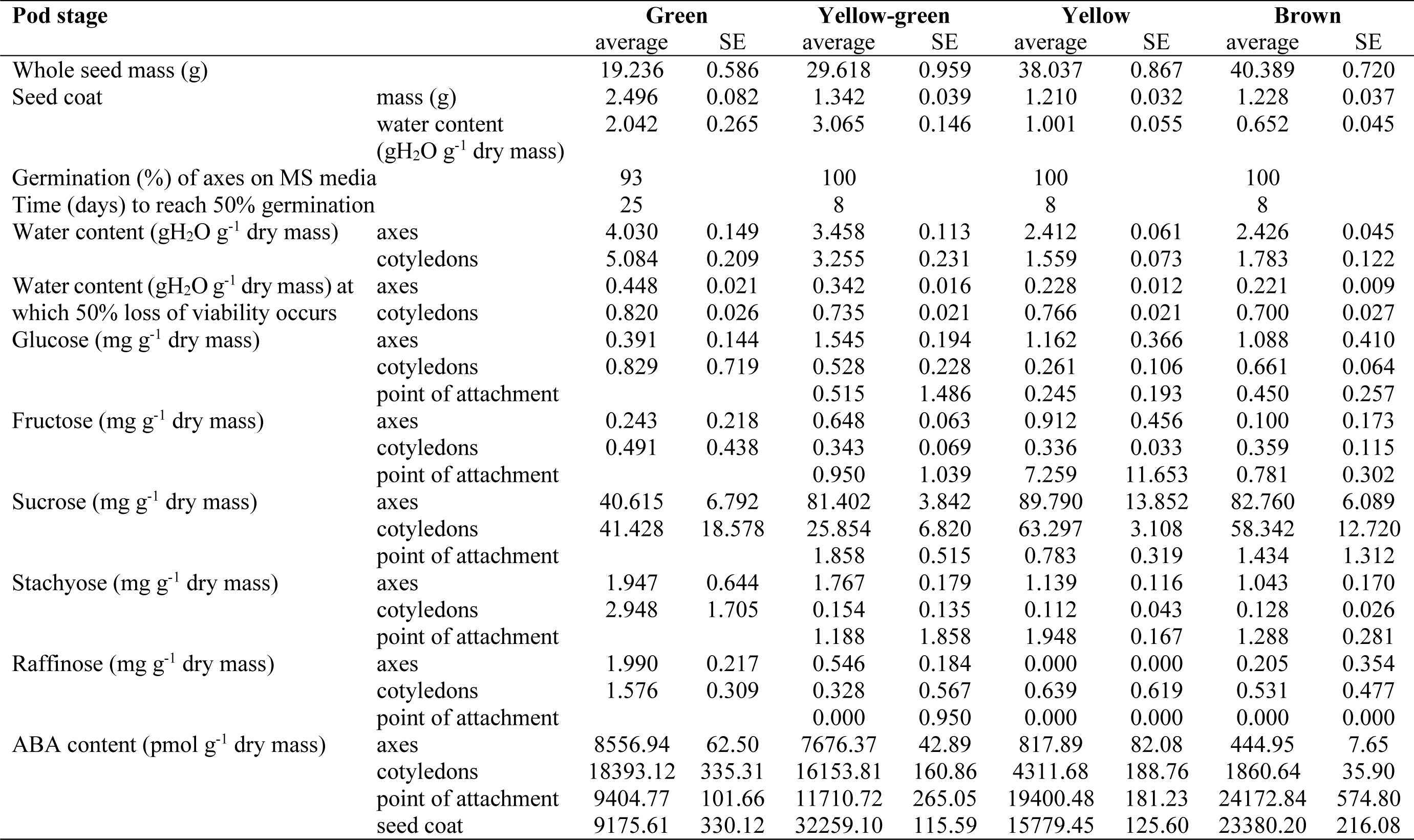
Phenotypic parameters associated with late seed development of *Castanospermum australe.* Point of attachment refers to tissue attaching the embryo to the seed pod.

While seeds had the capacity to germinate prior to reaching full embryonic size, germination rate was slow. Full germination capacity, typically of newly shed seeds was achieved at the yellow green pod stage (Table 1). The lethal water content of axes below which 50% viability was lost after drying declined from 0.45 gH_2_O/g^-1^ dry mass in those extracted from green pods, to 0.23 gH_2_O/g^-1^ dry mass in axes from yellow pods, after which there was no significant change. Cotyledons were more sensitive to dehydration, with 50% loss of viability occurring below 0.82 gH_2_O/g^-1^ dry mass in those from green pods, this declining slightly with development to 0.70 gH_2_O/g^-1^ dry mass in cotyledons from brown pods (Table 1).

ABA regulates many aspects of plant growth and development including embryo maturation, seed dormancy, germination, cell division and elongation. ABA content of *C. australe* embryos was high during early developmental stages but declined considerably in both axes and cotyledons in the transition from the yellow-green to the yellow pod stage (Table 1). ABA content increased considerably in the seed coat in the transition from the yellow to the brown pod stage, especially in the point of attachment (tissue that attaches the embryo to the pod).

### Seed maturation drying

DT seeds become tolerant of drying midway during seed development, concomitant with reserve accumulation (Chatelain *et al.*, 2012). From this stage onwards, there is progressive loss of water characterizing a process termed ‘maturation’ drying, that occurs after reserve accumulation is complete and full seed size is attained. There was no maturation drying typical of DT seeds after the yellow pod stage in *C. australe* (Table 1).

Previous work has identified transcripts that accumulate during the acquisition of desiccation tolerance in *M. truncatula* seeds (Terrasson *et al.*, 2013; Righetti *et al.*, 2015). Homologs of 121 of these genes failed to accumulate transcripts to a similar extent in *C. australe* cotyledons in comparable seed developmental stages (Table S1). These transcripts are related to sugar metabolism, photosynthesis, seed development, protection against abiotic stress and modulation of plant stress responses. Examples include *ABI3, ABI5*, chaperone proteins, heat shock factor proteins, putative LEAs, transparent testa protein, oleosins, *1-CYSTEINE PEROXIREDOXIN*, and α-galactosidases.

We identified 269 transcripts with decreasing abundance in *C. australe* during final maturation and increasing abundance in *M. truncatula* (Table S2). A certain number of these transcripts are possibly involved in longevity (life span in the dried state) (Verdier *et al.*, 2013; Righetti *et al.*, 2015). Some of these genes are related to metabolic and catabolic processes, such as *lipid metabolic process, cellular lipid metabolic process* and *nitrogen compound metabolic process* (Table S3), reflecting the type of reserves accumulated. Mature *C. australe* seeds predominantly accumulate starch (85% dry mass, Table1). Lipids and proteins constitute only 3-6% and 2.8% dry mass, respectively. Mature seeds of *M. truncatula* predominantly accumulate protein (30-40% dry mass), but also contain storage lipids (7-9% dry mass) and a small amount of starch (< 1% dry mass) (Djemel *et al.*, 2005).

We also identified 296 transcripts with decreasing abundance in *M. truncatula* and increasing abundance in *C. australe*. They are related to *root development, developmental process* and *regulation of localization* (Table S2), which are likely associated with germination related processes.

LEA proteins have been related to survival in the dry state (Chatelain *et al.*, 2012) and to responses to environmental stresses, including desiccation (Tunnacliffe & Wise, 2007). In *C. australe*, several LEA proteins failed to accumulate in cotyledons (Delahaie *et al.*, 2013). A genome-wide search for LEAs in the genome of *C. australe* identified 94 LEA motif-containing proteins, a number similar to what has been described for DT-seeded species, for example, 88 in *Arabidopsis thaliana* and 99 in *Sorghum* bicolor, as well as for vegetative DT in the resurrection plants *Oropetium thomaeum* and *Xerophyta viscosa* with 94 and 126 LEAs, respectively (VanBuren *et al.*, 2017; Costa *et al.*, 2017). A hierarchical clustering of expression of the LEAs in developing seeds of *C. australe* and *M. truncatula* separated the transcripts in two major clusters (Figure S1). LEAs in the first cluster belong to different families and transcript abundance increased considerably in *M. truncatula* seeds towards mid maturation. However, in *C. australe* most of these increased only slightly in early stages of development but declined during the later stages. LEAs in the second cluster belong to the LEA_2 family and do not undergo major changes in transcript abundance in either species. This family encodes ‘atypical’ LEA proteins because of their more hydrophobic character compared to other LEA families (Hundertmark & Hincha, 2008). Functional studies on LEA_2 proteins suggest that they do not act in the protection of membranes in tissues undergoing dehydration, although some proteins of this family were shown to have enzyme protective properties under both freezing and drying conditions (Dang *et al.*, 2014).

Changes in soluble sugar content and composition have been described as a characteristic of late maturation in DT seeds and correlate with the acquisition of longevity and preparation for the dry state (Wang *et al.*, 2013; Leprince *et al.*, 2017). Whereas in all DT legume seeds, raffinose family oligosaccharides (RFO) are the predominant sugars that increase during late maturation, *C. australe* seeds were composed of 7-10% of soluble sugars, with sucrose being the most abundant sugar detected and only minute amounts (0.7% of total soluble sugars at the brown pod stage) of RFOs accumulating. Low ratios of sucrose:RFO accumulation have been proposed to be a signature of desiccation tolerance and potential indicators of seed storage categories (Steadman *et al.*, 1996; Farrant *et al.*, 2012) and the opposite of this, as depicted in *C. australe* could be one of the reasons for desiccation sensitivity in this species.

### Genome sequencing and assembly

In an effort to obtain a genomic blueprint of DS seeds we sequenced the genome of *C. australe*. This species has a key position in the legume family phylogenetic tree, in the basis of the ADA clade, which favours the study of trait evolution as well as the ancient polyploid history of papilionoid legumes.

We produced an assembly with a total length of 382 Mb and an N50 of 832.6 Kb that covers 96.7% of the predicted genome size. The assembly consisted of 1,210 contigs and 1,027 scaffolds (Table 2). The GC content was 32.9%. Genome annotation identified 29,124 protein-coding genes of which 98.1% show high sequence similarity to proteins in TrEMBL and 84.4% in Swiss-Prot databases. An estimation of genome completeness indicated that 96.4% of the BUSCO (Benchmarking Universal Single-Copy Orthologs) genes were present. Transposable elements covered 15.5% of the total genome. Repeat elements comprised 119 Kb of SINE (Short Interspersed Nuclear Element) retrotransposons, 383 Kb of LINE (Long Interspersed Elements) retrotransposons, 13 Mb DNA transposons and 42 Mb of annotated LTR (long terminal repeat-retrotransposons) sequences (Table 2).

**Table 2.**
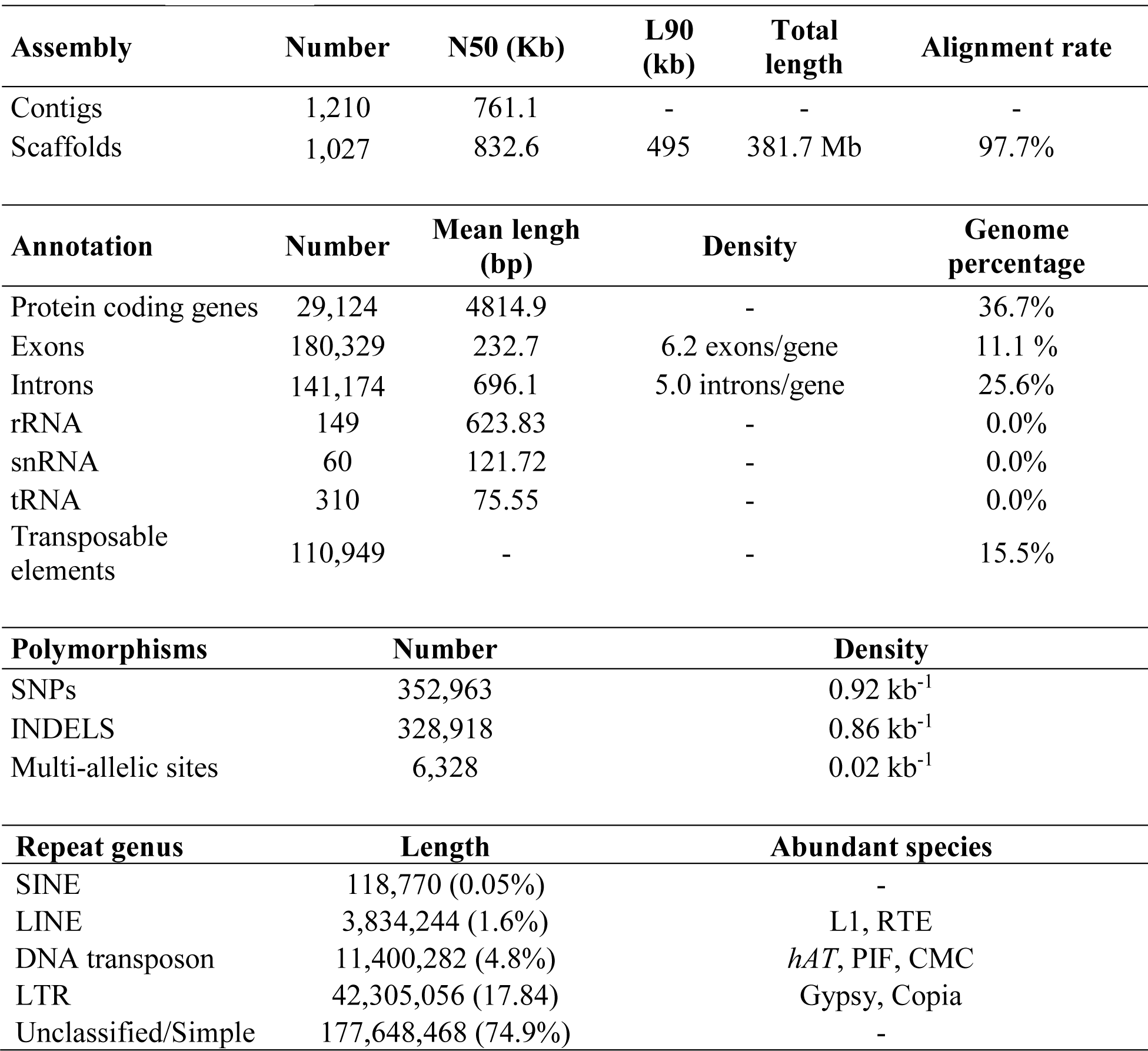
Overview of assembly, annotation, polymorphism and repeat elements on the *Castanospermum australe* genome. N50: scaffold size above which 50% of the total length of the sequence assembly can be found. L90: number of contigs whose summed length contains at least 90% of the sum of the total length of the sequence assembly. rRNA: ribosomal RNA. snRNA: small nuclear RNA. tRNA: transfer RNA. SNP: single nucleotide polymorphism. INDEL: insertion or deletion of bases in the DNA. SINE: short interspersed nuclear element. LINE: long interspersed nuclear element. LTR: long terminal repeat. L1: LINE-1. RTE: retrotransposable element. *hAT*: *hobo/Ac/Tam3*. PIF: P Instability Factor. CMC: CACTA/Mirage/Chapaev.

### Genomic alterations linked to seed desiccation sensitivity

To investigate the evolution of *C. australe* and legume diversification, a phylome was constructed. A phylome constitutes the collection of all gene phylogenies in a genome. It is a valuable source of information to establish evolutionary relationships among organisms and their genes (Huerta-Cepas *et al.*, 2014). Our phylome contained the evolutionary histories of all *C. australe* protein coding genes and their homologues in 20 publicly available sequenced plant species (Figure 2). These species represent a broad phylogenetic distribution and include the diverse seed storage phenotypes, i.e. DT, DS and intermediate seeds. Intermediate seeds are typically tolerant of relatively extreme water loss, to 0.1-0.14 g H2O/g^-1^ dry mass but no lower than this(Marques *et al.*, 2018), and have poor survival under conventional storage conditions (Berjak & Pammenter, 2013). Such seeds thus show a storage phenotype in-between orthodox and recalcitrant seeds.

The phylome analysis indicated that very few protein-coding genes (≤ 1%) were present in DS species only, and no protein-coding genes were retained in all DT species but lost in all DS species (Figure 2). Several genes were lost in all DS species and retained in at least half of the DT species (Table S4). 76 genes lost their ortholog but kept a paralog in *C. australe* and 59 genes were lost in *C. australe* without detected paralogs, of which 11 were shared with other DS-seeded species.

According to the phylome, 3,716 gene families were expanded exclusively in *C. australe*, of which 180 were associated with transposons. Expansion size ranged from 2 to 32 genes and involved 30.5% of the predicted proteome. After removal of expansions associated with transposable elements or viruses, the remaining expanded gene families were enriched for GO (gene ontology) terms such as *defence response, flavonoid biosynthetic process, terpene synthase activity* and *nutrient reservoir activity*. GO terms associated with *terpene synthase activity, lyase activity* and *pectinesterase activity* were also enriched at the base of the Papilionoideae subfamily. The phylome analysis also indicated that the duplication frequency at the base of the Papilionoideae subfamily is lower (0.22) than that found at the base of the Fabaceae family (0.73, Figure 3). Histograms of the synonymous rates and average rates of syntenic blocks for six legume species showed distinctive peaks tracing back to the shared papilionoid legume whole genome duplication (WGD), suggesting that the rate of evolution is very diverse among closely related family members (Figure 4). The peak in *C. australe* corresponds to a rate of synonymous (Ks) site mutations of 0.25 which is less than half of the rate observed in *G. max* (0.6) and a third of that in *M. truncatula* (0.85). The substitution rate in *C. australe* is so low that a second peak corresponding to the eudicot hexaploidy (gamma WGD event) is still visible. The gamma event was also detected in the histograms of block averages in *G. max* and *P. vulgaris*. The low Ks rate in *C. australe* is likely due to this species being the only perennial in this comparative study and the one with the longest generation times.

**Figure 3.**
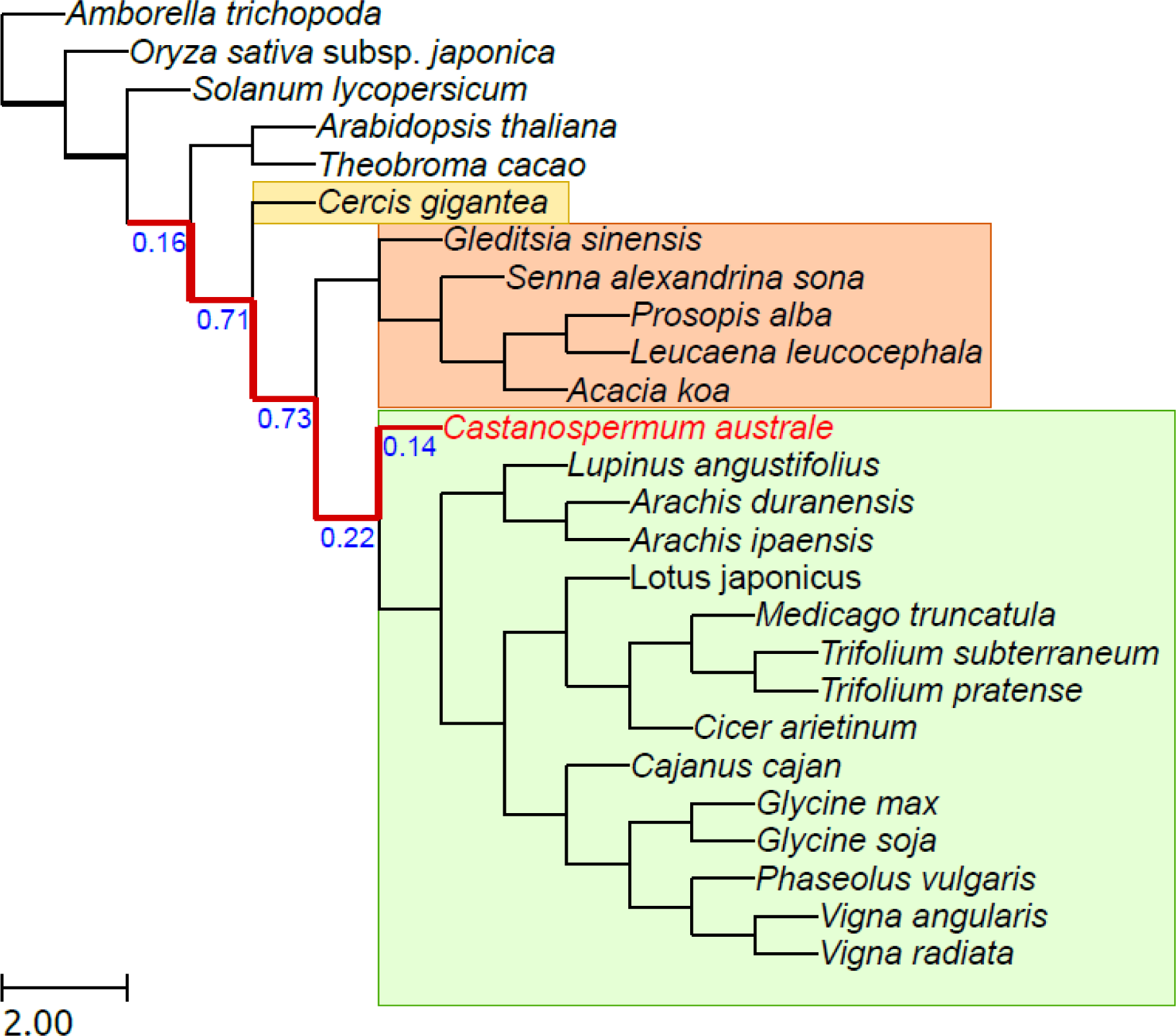
Phylogenetic tree showing duplication rates of selected species including species of the subfamilies: Mimosoideae (inside yellow rectangle) and Caesalpinioideae (inside orange rectangle) and Papilionoideae (inside green rectangle). Phylogentic tree based on: Lavin, M., Herendeen, P.S. and Wojciechowski, M.F. (2005) evolutionary rates analysis of leguminosae implicates a rapid diversification of lineages during the tertiary. Systematic Biology 54, 575–594.

**Figure 4.**
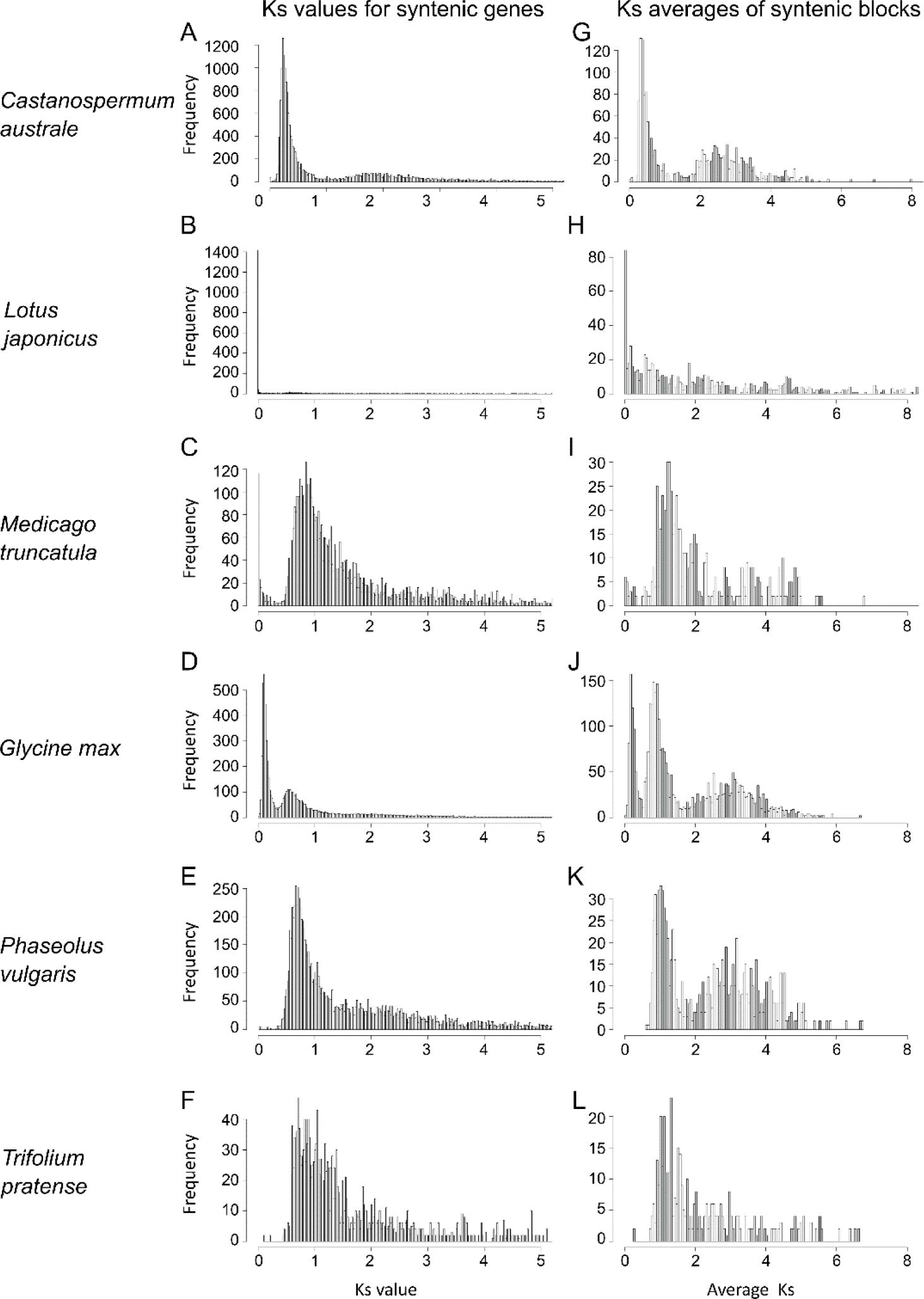
Synonymous mutations of duplicated genes and syntenic blocks. Histograms of synonymous mutations (Ks) of duplicated genes (A-F) and average Ks of syntenic blocks (G-L). (A-B) *Castanospermum australe*, (C-D) *Lotus japonicus*, (E-F) *Medicago truncatula*, (G-H) *Glycine max*, (I-J) *Phaseolus vulgaris*, (K-L) *Trifolium pratense.*

The evolution of a trait is shaped by the selective pressures to which it is subject. Some selective pressures act to increase the benefits accumulated while others act to reduce the costs incurred, affecting the cost/benefit ratio. Different selective pressures can be estimated by the ratio of the number of nonsynonymous substitutions per non-synonymous site (dN) in a specific period to the number of synonymous substitutions per synonymous site (dS) in the same period (Mugal *et al.*, 2014).

Genome wide analysis of protein coding genes of *C. australe* in comparison with other legume genomes enabled the identification of genes with 2-fold higher dN/dS in *C. australe* (Table S5). Among these were genes associated with hormonal signalling, such as *ACTIVATION-TAGGED BRI1* (*BRASSINOSTEROID-INSENSITIVE1*)-*SUPPRESSOR1* (*ATBS1*), Ethylene Insensitive 3 family protein, *BRI1-ASSOCIATED RECEPTOR KINASE* (*BAK1*) and *SALT TOLERANCE HOMOLOG2* (*STH2*). Other examples are: *SEEDSTICK* (*STK*), *TRANSPARENT TESTA5* (*TT5*), *ENDO-BETA-MANNANASE7* (*MAN7), FLOWER FLAVONOID TRANSPORTER* (*FFT*) and *ROTUNDIFOLIA3* (*ROT3*).

The evolution of a trait can also be investigated by analysing the degree to which genes remain on corresponding chromosome (synteny) and in corresponding orders over time. We investigated whether the loss of synteny in *C. australe* genes could be related to the loss of seed desiccation tolerance. There are 169 genes re-arranged in the *C. australe* genome that have syntenic orthologs in 50 angiosperm species. Most noteworthy among these were the genes *BRASSINOSTEROID INSENSITIVE 3* and *5* (*BIN3* and *BIN5*), which participate in brassinosteroid (BR) signalling and are associated with seed size (Yin *et al.*, 2002). Furthermore, genes such as *MINISEED3* (*MINI3*) and *HAIKU1* (*IKU1*), regulators of seed size via the BR pathway in *A. thaliana* (Luo *et al.*, 2005), also lost synteny in *C. austral*e, which could contribute to the large seed size in this species.

The synteny between the genome of *C. australe* and other legume species was evaluated by aligning the genome of *C. australe* against itself and against the genomes of *G. max* and *M. truncatula* (Figure S2). Most of the *C. australe* genome is syntenic with itself and mostly duplicated after the WGD event. While the duplicates are associated with the most recent WGD event, many paralogs derived from the gamma event were also detected. The alignment of *C. australe* against *G. max* indicated a high amount of syntenic orthologs and paralogs, whereas the alignment of *C. australe* against *M. truncatula* indicated that although many syntenic orthologs have been conserved, most of the WGD-derived paralogs were lost. Moreover, most of the duplicated regions retained by *M. truncatula* were also duplicated and retained in *C. australe*.

## Discussion

In orthodox seeds, survival in the dry state is a result of a series of molecular and cellular processes that occur during the late stages of seed development. These processes result in the acquisition of desiccation tolerance and longevity in the dry state. Although we have a detailed understanding of these associated processes in orthodox seeds, limited information is available regarding the development of seeds that do not fully activate them, such as *C. australe*. Our study provides detailed information about *C. australe* seed development and desiccation sensitivity.

In *C. australe* axes and cotyledons, some water loss occurss during reserve accumulation but this stops once the embryo reaches its full size, stabilizing at 2.4 and 1.6 gH_2_O g^-1^ dry mass, respectively (Table 1). This value is substantially higher than the 0.1 gH_2_Og^-1^ dry mass reached by desiccation-tolerant seeds, such as *M. truncatula*.

The pattern of sugar accumulation also differs markedly between these species. The percentage of soluble sugars in seeds of *C. australe* (7-10%) is comparable with the average percentage for legume species (8-10% (Djemel *et al.*, 2005)). However, while RFOs are the main sugars in *M. truncatula*, comprising 90% of the total soluble sugar content, in *C. australe* only minute amounts of RFOs, mainly stachyose, could be detected (0.7% of total soluble sugars at the brown pod stage). Stachyose and raffinose contents were highest in seeds from the green pod stage and decreased with further progress of maturation (Table 1). A similar finding has been reported for the non-viviparous highly DS seeds of *Avicennia marina* (Farrant *et al.*, 1992). In parallel, sucrose and glucose content increased. The reduction of stachyose content during further maturation suggests hydrolysis, normally occurring in germinating DT seeds (Rosnoblet *et al.*, 2007). A comparative analysis of transcripts linked to RFO metabolism between *C. australe* and *M. truncatula* identified transcripts of genes related to the synthesis of sucrose from fructose-6 phosphate that remained high in *C. australe* whereas they decreased in *M. truncatula*. Conversely, the transcripts of several genes related to the synthesis of galactinol or raffinose and stachyose accumulated during development of *M. truncatula* seeds while their abundance remained low in *C. australe* (Supplementary Figure S1B). This set of genes might explain the lack of RFO accumulation in *C. australe* seeds. Whereas the specific roles of RFOs in protection compared to the nonreducing sucrose remain unconfirmed, the DS *A. thaliana abi3* mutants as well as Mt-*abi5* are also impaired in the accumulation of RFOs (Zinsmeister *et al.*, 2016).

At the transcriptome level, transcripts with decreasing abundance in *M. truncatula* and increasing abundance in *C. australe* reinforce the notion that towards the end of seed development, *C. australe* is metabolically active while *M. truncatula* is entering a phase of low metabolic activity and quiescence. Examples of these transcripts are beta-galactosidase (Medtr8g039160), xyloglucan galactosyltransferase (Medtr1g069460) and TCP family TF (Medtr6g015350). Interestingly, these genes lost synteny in *C. australe* compared to their *M. truncatula* orthologs. Their involvement in carbohydrate metabolism and control of cell proliferation hints at implication in the germination program. The germination program remains active in *C. australe*, as DS seeds generally do not display developmental arrest, allowing the maintenance of high metabolic activity. In contrast, transcripts of indole-3-acetic acid-amido synthetases, involved in auxin homeostasis, accumulated in *M. truncatula* during development but decreased in *C. australe*. Auxin has been reported to maintain seed dormancy by interacting with ABA (Liu *et al.*, 2013).

At the genome level, very few protein-coding genes (≤ 1%) were present in DS species only (Figure 2), supporting the hypothesis that independent evolutionary events gave rise to DS-seeded species. No protein-coding genes were retained in all DT species and lost in all DS species. However, several were lost in all DS and retained in at least half of the DT species. Among these were the transcription factors (TFs) *VERDANDI* and *MYB44*-*like*. *VERDANDI* participates in ovule identity complex and, when mutated, affects embryo sac differentiation in *A. thaliana* (Matias-Hernandez *et al.*, 2010; Mendes *et al.*, 2016). Interestingly, *VERDANDI* and *MYB44*-*like* were also retained in intermediate-seeded species.

One gene (*PLAC8*) was lost without retention of paralogs in three out of four DS-seeded species, namely *C. australe, Castanea mollissima* and *Elaeis guineensis*. The knock-out of this gene caused increased seed and fruit size in maize (Libault & Stacey, 2010). In addition, the fw2.2 locus containing the *PLAC8* gene has been suggested to be the key to the evolution of tomato fruit size (Frary, 2000). Large seeds and fruits are common features of DS species and presumably reduce the rate of seed drying and hence the risk of desiccation-induced embryo mortality (Daws *et al.*, 2006).

Amongst the genes that lost synteny in *C. australe* without detected paralogs, 11 were shared with other DS-seeded species. Two of these genes, *LEA2* and *FIBRILLIN5* accumulate transcripts in *M. truncatula* during seed maturation and upon re-induction of desiccation tolerance in germinated seeds (Terrasson *et al.*, 2013). The gene *GUN5*, a magnesium chelatase involved in retrograde signalling and ABA signalling to the nucleus (Jiang *et al.*, 2014), was lost in *C. australe* without paralogs. This pathway is affected in *M. truncatula abi5* mutants that produce seeds with strongly reduced longevity (Zinsmeister *et al.*, 2016) and cannot reacquire desiccation tolerance after germination (Terrasson *et al.*, 2013).

Examples of genes which lost their ortholog but kept a paralog in *C. australe* are *RETARDED ROOT GROWTH-LIKE* (*RRL)* and *MOTHER OF FT* (*MFT*), involved in ABA- and BR-signalling. *RRL* mediates ABA signal transduction through *ABI4* (Park *et al.*, 2015) and *MFT* regulates seed germination and fertility involving ABA- and BR-signalling pathways (Sun *et al.*, 2010).

ABA is involved in the formation of mature DT seeds, and inhibition of their subsequent germination under conditions unfavourable for seedling growth (Finkelstein, 2013). During seed development, an increase in ABA content has been related to a transition from growth by cell division to growth by cell enlargement and to cell cycle arrest at the G1/S transition. This increase may be related to the role of ABA in promoting senescence, a process which precedes abscission (Finkelstein, 2013). Additionally, the higher ABA content in the seed coat could play a role in delaying germination of the embryo until ideal conditions for germination are met, or to aid temporal and/or spatial dispersal without immediate loss of viability. Interestingly, the seed coat starts to peel away from the embryo in mature seeds (Figure 1) and this increases markedly once seeds are shed, suggesting increasing lack of inhibition of germination of the embryo.

ABA is also a key regulator of abiotic stress responses and acquisition of desiccation tolerance during seed development (reviewed by (Dekkers *et al.*, 2015)). Disruption of ABA biosynthesis or -signalling leading to lack of or insensitivity to ABA results in loss of seed desiccation tolerance (Verdier *et al.*, 2013). We observed alterations in genes related to ABA signalling, such as *MFT, GPCR-TYPE G PROTEIN 1* (*GTG1*), *STH2, ABI3* and *ABI5. C. australe* lost an ortholog of *MFT* but maintained a paralog. Mutations in this gene may cause ABA hypersensitivity at germination and is associated with dormancy (Vaistij *et al.*, 2013). *STH2* is involved in ABA signalling, is highly expressed during embryogenesis (Xu *et al.*, 2014) and has a high dN/dS in *C. australe. GTG1* was lost in *C. mollissima* and its knock-out causes ABA hyposensitivity in *A. thaliana* seeds (Pandey *et al.*, 2009). *ABI3* and *ABI5* showed contrasting expression patterns during *C. australe* seed development compared to *M. truncatula*. These two TFs have been shown to play essential roles in seed development and acquisition of desiccation tolerance (Terrasson *et al.*, 2013; Dekkers *et al.*, 2015; Zinsmeister *et al.*, 2016).

BRs have been implicated in seed development and are known to antagonize seed dormancy and stimulate germination (Steber, 2001). In *C. australe,* several genes involved in BR-biosynthesis and -signalling and seed development have undergone genetic changes (Figure 5). For example, *BRASSINOSTEROID INSENSITIVE-LIKE 3* (*BRL3*) lost an ortholog but kept a paralog; *IKU1* and *MINI3* lost synteny; and *ROT3, DE-ETIOLATED 2* (*DET2*), *ATBS1* and *BAK1* have higher dN/dS in *C. australe* than in *M. truncatula.* Furthermore, an ortholog of *ATBS1* was highly expressed during late development of *C. australe* contrasting with the decreasing expression in *M. truncatula* seeds. These data support the hypothesis that subcellular metabolism associated with germination is initiated during the late stages of development in the DS seeds of *C. australe*. Overall, our results support the hypothesis that the evolution of desiccation sensitivity was not caused by massive alterations in enzymes and structural proteins but instead by discrete mutations in regulatory genes.

**Figure 5.**
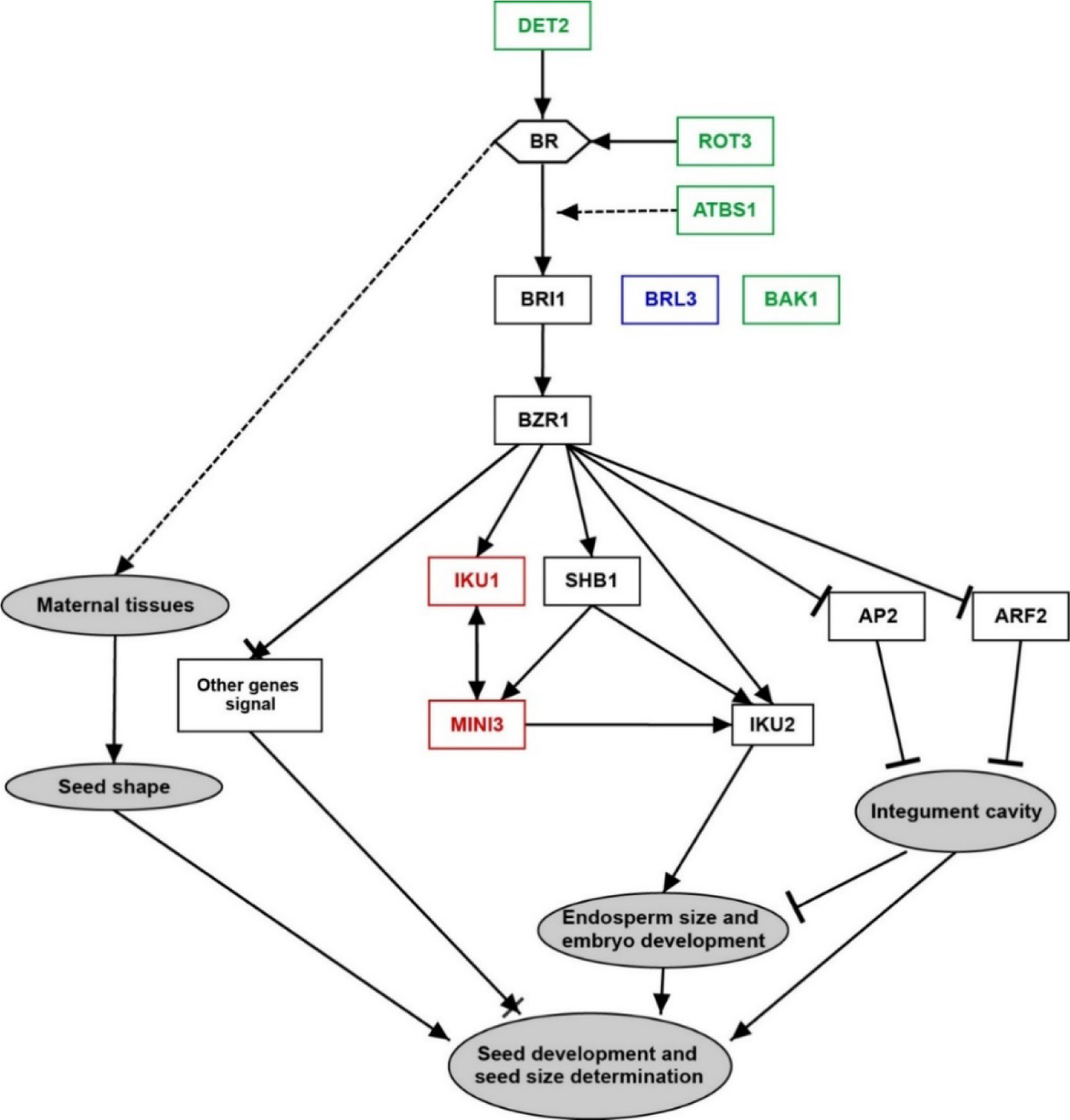
Hypothetical model for brassinosteroid-regulated seed development (adapted from Jiang et al. (2013)). Red shapes indicate genes that lost synteny in *Castanospermum australe* compared to *Medicago truncatula*. Green shapes indicate genes with higher dN/dS in *C. australe* than in *M. truncatula*. Blue shapes indicate genes that lost an ortholog, but kept a paralog in *C. australe*. BR: brassinosteroid. ATBSI1: *ACTIVATION-TAGGED BRI1 (BRASSINOSTEROID-INSENSITIVE1)-SUPPRESSOR1*. AP2: *APETALA 2*. ARF2: *AUXIN RESPONSE FACTOR 2*. BAK1: *BRI1-ASSOCIATED RECEPTOR KINASE 1.* BRI1: *BRASSINOSTEROID INSENSITIVE 1*. BRL3: *BRASSINOSTEROID INSENSITIVE-LIKE 3*. BZR1: *BRASSINAZOLE RESISTANT 1*. DET2: *DE-ETIOLATED 2*. IKU: *HAIKU*. MINI3: *MINISEED 3*. ROT3: *ROTUNDIFOLIA 3*. SHB1: *SHOEBOX 1*.

Natural populations often undergo the weakening or removal of a selective force that had been important in the maintenance of a trait, characterizing a “relaxed selection” (Lahti *et al.*, 2009). When a DT-seeded species is subjected to an environment where desiccation tolerance is not an adaptive trait, there should be relaxation of its evolutionary constraints that can eventually lead to its loss. DS-seeded species evolved in environments where the conditions favour immediate germination and seeds are programmed to initiate germination upon, or shortly after shedding (Farrant *et al.*, 1993a; Daws *et al.*, 2006; Berjak & Pammenter, 2013). DT seeds, which very often display a form of dormancy, normally form seed banks in the soil. In contrast, DS seeds germinate immediately and usually form seedling banks under shaded forest canopy and take advantage of an eventual light gap for faster establishment. Furthermore, in these species, the generally increased seed size favours seedling establishment under shaded forest conditions (Daws *et al.*, 2006).

In summary, seed desiccation sensitivity evolved multiple independent times in environments where water is highly abundant and predictable across long periods, favouring immediate seed germination. In such environments, the evolutionary pressure for DT seeds is relaxed and the production of DS seeds is not disadvantageous. This was the case for *C. australe*. Among the currently known DS-non-viviparous seeds, *C. australe* is one of the most sensitive to water loss. We have pinpointed some of the factors behind this sensitivity, namely displacements, loss of synteny and mis-expression of specific genes related to the BR- and ABA-signalling pathways, carbon metabolism, control of cell proliferation, protection against abiotic stresses and modulation of plant stress responses. These alterations are likely to have led to an increased seed size; high starch and low protein and lipid seed content; low accumulation of LEA proteins and RFOs in the seeds; and failure to start the seed maturation drying phase.

The low similarity between DS-seeded species confirms the hypothesis that desiccation sensitivity evolved independently. Moreover, it supports the idea that although the evolution of many factors was necessary for the appearance of seed desiccation tolerance, only a few changes in some of these factors are enough for its loss.

## Author contributions

A.M. and M.-C.D.C. wrote the article; U.C. performed physiological experiments; M.-C.D.C., E.J., T.Z., M.D. and H.N. performed the bioinformatics; A.M., E.S., M.M.-H., I.J. and T.G. contributed to the genome and transcriptome analysis; J.D. and J.B. performed and analysed the transcriptomics; M.E.S. performed the PacBio sequencing and initial genome analysis; J.B., H.W.M.H. and J.M.F. initiated and coordinated the work and directed preparation of the article.

## Supporting information

All supplemental information

## Acknowledgements

A.M. received financial support from CNPq–National Council for Scientific and Technological Development (246220/2012-0) Brazil. J.M.F. contributed towards this work from funding from the National Research Foundation (grant number 69416) and her DST-NRF South African Research chair (grant number 98406). This work was funded in part by a grant from the Region des Pays de la Loire, France (QUALISEM 2009-2013) and the bilateral Partenariat Hubert Curien (PHC) program France–South Africa (grant no. 25903RE) to O.L. and J.B.). We acknowledge David Lalanne and the ANAN platform of the SFR Quasav, Angers, France for the assistance with the microarray analysis. We acknowledge Bas te Lintel Hekkert for library preparation for genome sequencing.

## Supporting Information

**Figure S1.** Hierarchical clustering of expression values.

**Figure S2.** Phylogenetic tree showing duplication rates of selected species.

**Table S1**: Gene ontology (GO) enrichment analysis of biological processes in relation to the acquisition of tolerance to water loss and to seed maturation in *Medicago truncatula* and *Castanospermum australe*.

**Table S2.** Genes changing transcript abundance in *Castanospermum australe* and *Medicago truncatula* in comparable seed developmental stages.

**Table S3.** Genes changing transcript abundance in *Castanospermum australe* and *Medicago truncatula* in comparable seed developmental stages during final maturation.

**Table S4.** Protein-coding genes lost in all DS and retained in at least half of the DT species.

**Table S5.** *Castanospermum australe* protein-coding genes with dN/dS (number of nonsynonymous substitutions per non-synonymous site (dN) in a given period of time divided by the number of synonymous substitutions per synonymous site (dS) in the same period) ratio ≥2.

