## Supplementary material for "A blueprint of seed desiccation sensitivity in the genome of *Castanospermum australe*": All supplemental information

The following Supporting Information is available for this article:

Article acceptance date: Click here to enter a date.

**Fig. S1.** Hierarchical clustering of expression values. (A) Late embryogenesis abundant (LEA)-domain containing genes, (B) genes involved in RFO synthesis in developing seeds of *C. australe* and *Medicago truncatula*. C2.5: cotyledons weighing between 1.51 and 2.5g. C4.5: cotyledons weighing between 3.51 and 4.5g. C7.5: cotyledons weighing between 5.51 and 7.5g. GC: green cotyledons. YGC: yellow-green cotyledons. BC: brown cotyledons. GA: green axes. YGA: yellow green axes. BA: brown axes. DAP: days after pollination. ABS: pod abscission. DS: dry seeds. SMP: seed maturation proteins.

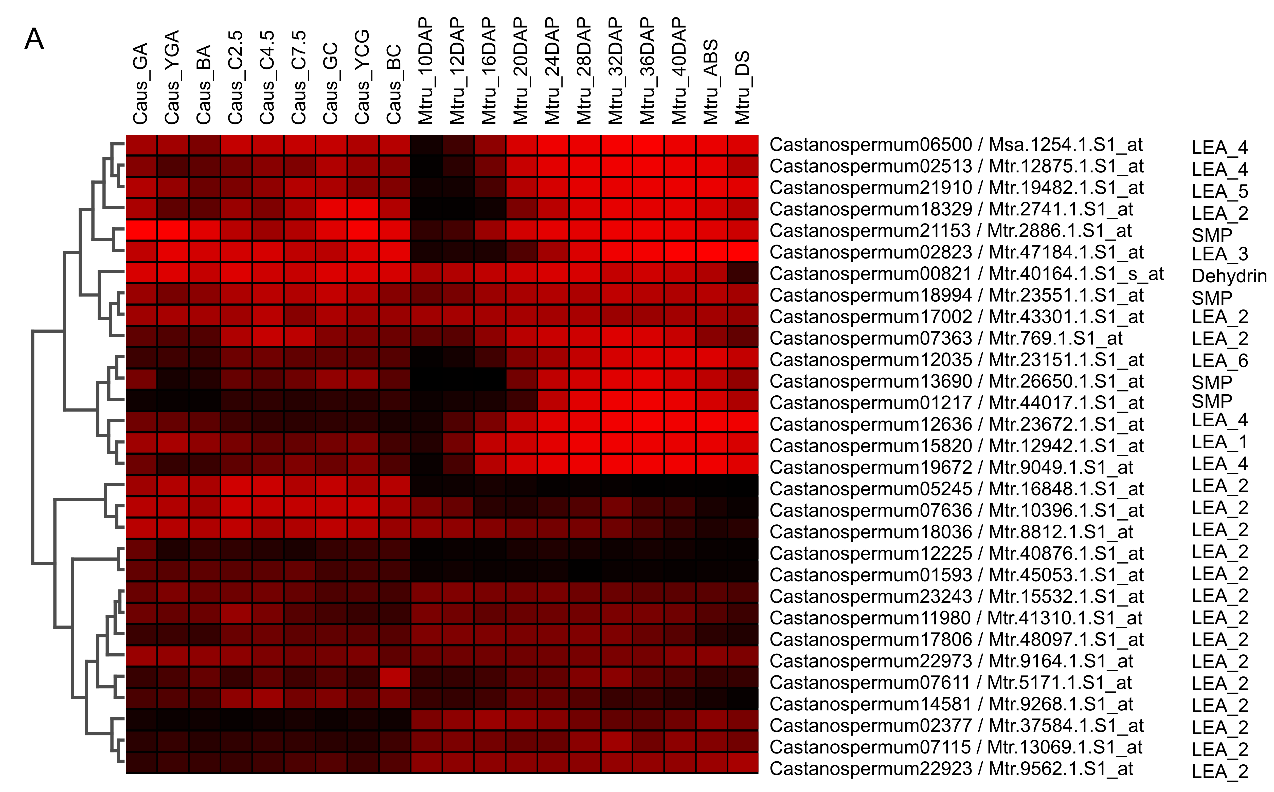

**B**

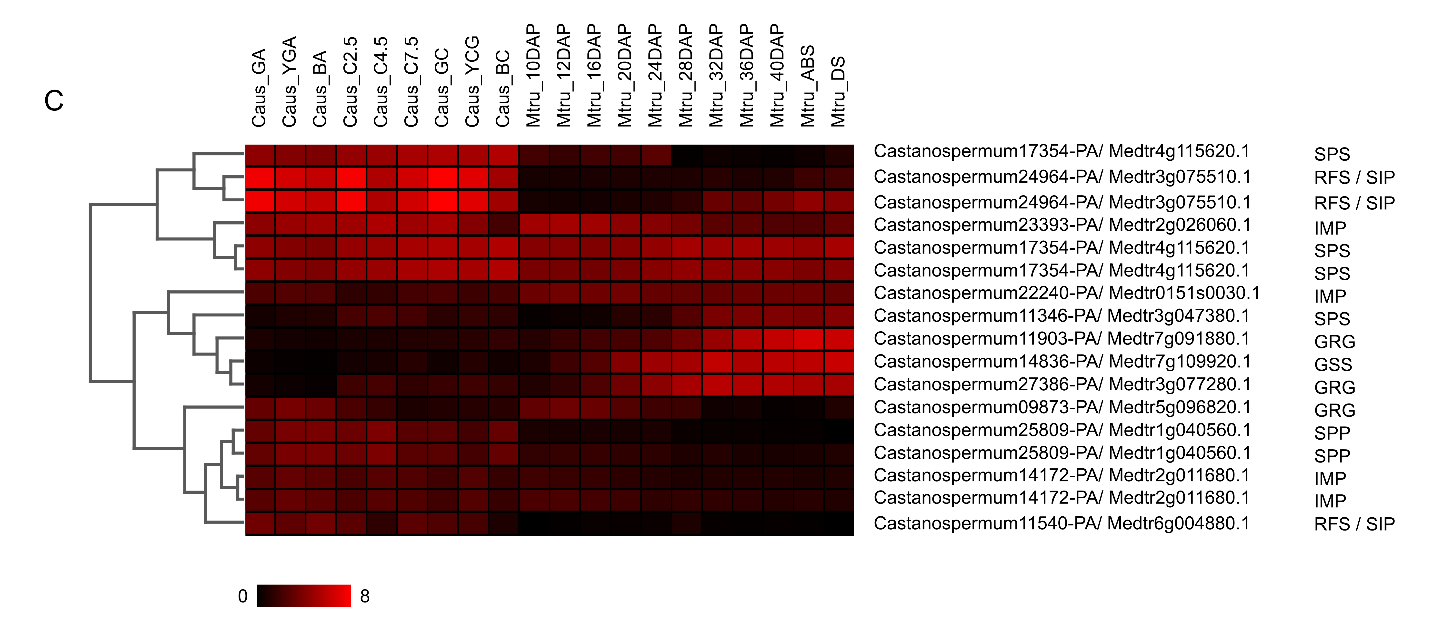

**Fig. S2.** (A-C) Histograms of log10 transformed Ks (rate of synonymous mutations) of syntenic gene pairs. These were identified through the alignment of (A) *Castanospermum australe* with itself, (B) *C. australe* with *Medicago truncatula*, and (C) *C. australe* with *Glycine max*. (D-F) Syntenic dotplots of comparisons of (D) *C. australe* with itself, (E) *C. australe* with *M. truncatula*, and (F) *C. australe* with *G. max*. Syntenic dotplots are coloured by their Ks values shown in A-C.

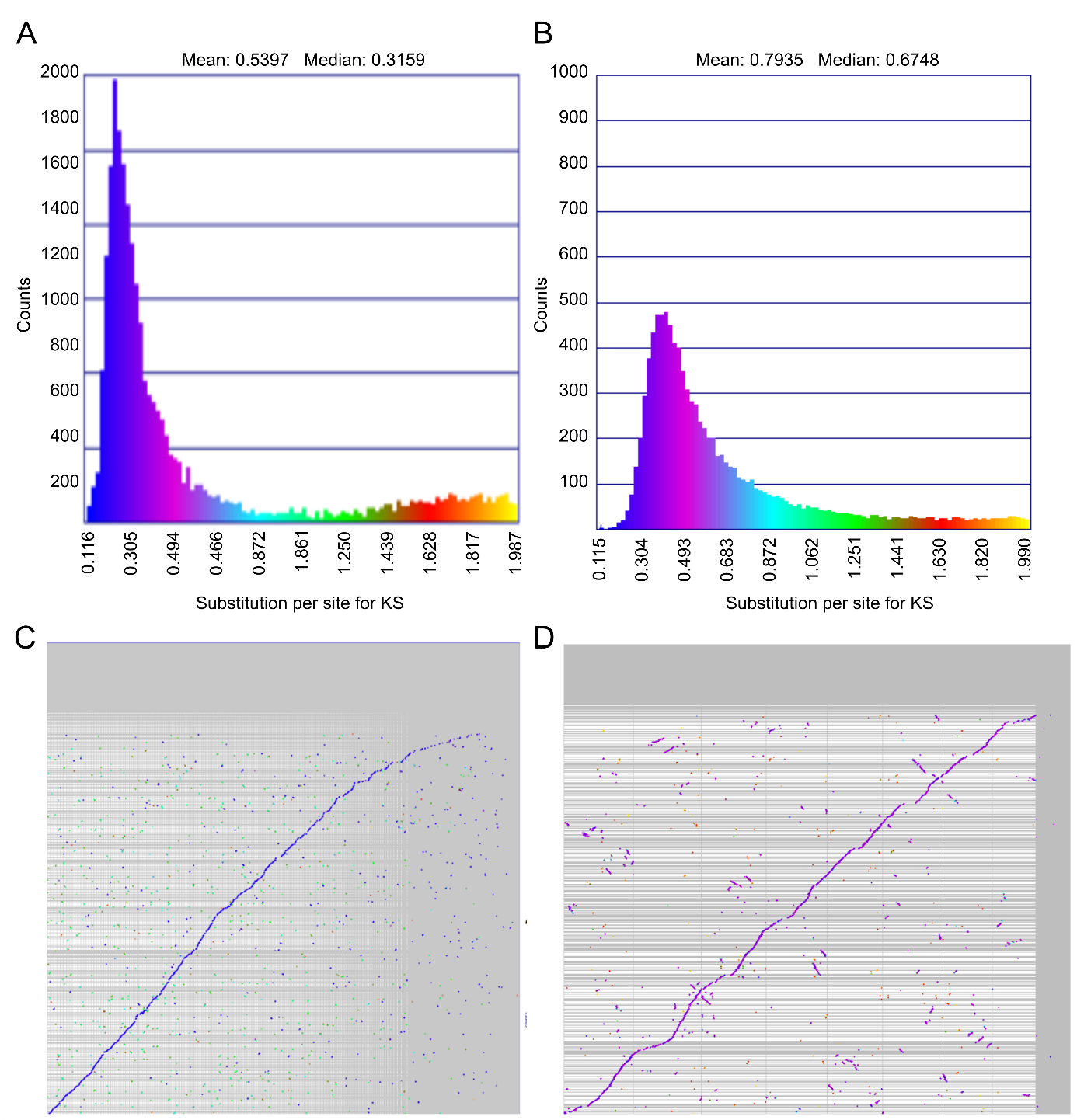

**Table S1.** Genes changing transcript abundance in *Castanospermum australe* and *Medicago truncatula* in comparable seed developmental stages. *C. australe* transcripts significantly changing abundance in the green pod stage compared to the stage when cotyledons weights between 1.51 and 2.50g were used. *M. truncatula* transcripts in the “desiccation tolerance module” identified by Righetti et al. (2015) or increasing abundance during the re-induction of desiccation tolerance in germinated seeds were used (Terrasson et al. 2013). dec: genes decreasing transcript abundance. inc: genes increasing transcript abundance. nc/nd: genes either not changing or not detected in the microarrays.

**See separate Excel file Table S1.xlsx**

**Table S2.** Genes changing transcript abundance in *Castanospermum australe* and *Medicago truncatula* in comparable seed developmental stages during final maturation. *C. australe* transcripts significantly changing abundance in the green pod stage (GC) compared to the brown pod stage (BC) were used. *M. truncatula* transcripts changing abundance in dry seeds (DS) compared to seeds 28 days after pollination (28DAP) were used (Verdier et al. 2013). dec: genes decreasing transcript abundance. inc: genes increasing transcript abundance. nc/nd: genes either not changing or not detected in the microarrays.

**See separate Excel file Table S2.xlsx**

**Table S3.** Gene ontology (GO) enrichment analysis of biological processes in relation to the acquisition of DT and to seed maturation in *Medicago truncatula* and *Castanospermum australe*. Genes increasing transcript abundance in M. truncatula during the phase of acquisition of tolerance to water loss were compared to genes decreasing transcript abundance in C. australe in the stage when the cotyledons weight between 1.51 and 2.50g in relation to green pod stage. Genes changing transcript abundance in M. truncatula at 28 or 32 days after pollination (DAP) in relation to dry seeds (DS) were compared to genes changing transcript abundance in C. australe cotyledons at the green pod stage (GC) in relation to cotyledons at the brown pod stage (BC). P-values of the false discovery rate (FDR) are shown together with the number of genes (#) in each GO term. Analysis was performed using AgriGO with the Arabidopsis TAIR10 background.

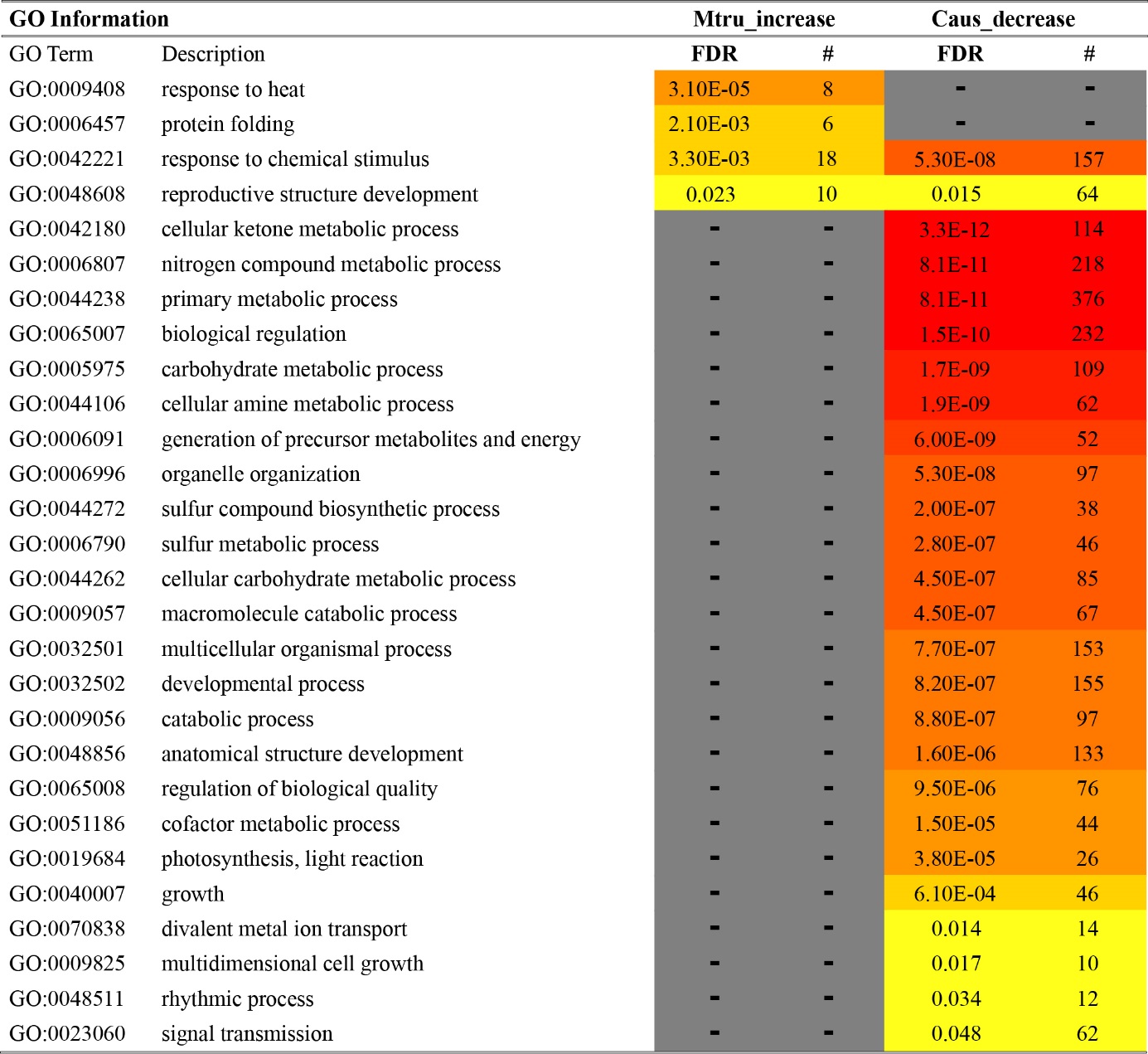

**Table S4**. Protein-coding genes lost in all DS and retained in at least half of the DT species.

**See separate Excel file Table S4.xlsx**

**Table S5**. *Castanospermum australe* protein-coding genes with dN/dS (number of nonsynonymous substitutions per non-synonymous site (dN) in a given period of time divided by the number of synonymous substitutions per synonymous site (dS) in the same period) ratio ≥2. Ratio was calculated by dividing C. australe dN/dS by the dN/dS average of other legume species. Caus: Castanospermum autrale. Gmax: Glycine max. Gsoj: Glycine soja. Pvul: Phaseolus vulgaris. Tpra: Trifolium pratense

**See separate Excel file Table S5.xlsx**
